# Bacterial cell size modulation along the growth curve across nutrient conditions

**DOI:** 10.1101/2024.09.24.614723

**Authors:** César Nieto, Claudia Igler, Abhyudai Singh

**Author notes:** Authors contributed equally (César Nieto, Claudia Igler).

## Abstract

Under stable growth conditions, bacteria maintain cell size homeostasis through coordinated elongation and division. However, fluctuations in nutrient availability result in dynamic regulation of the target cell size. Using microscopy imaging and mathematical modelling, we examine how bacterial cell volume changes over the growth curve in response to nutrient conditions. We find that two rod-shaped bacteria, *Escherichia coli* and *Salmonella enterica*, exhibit similar cell volume distributions in stationary phase cultures irrespective of growth media. Cell resuspension in rich media results in a transient peak with a five-fold increase in cell volume ≈ 2h after resuspension. This maximum cell volume, which depends on nutrient composition, subsequently decreases to the stationary phase cell size. Continuous nutrient supply sustains the maximum volume. In poor nutrient conditions, cell volume shows minimal changes over the growth curve, but a markedly decreased cell width compared to other conditions. The observed cell volume dynamics translate into non-monotonic dynamics in the ratio between biomass (optical density) and cell number (colony-forming units), highlighting their non-linear relationship. Our findings support a heuristic model comparing modulation of cell division relative to growth across nutrient conditions and providing novel insight into the mechanisms of cell size control under dynamic environmental conditions.

## Introduction

Bacteria modulate their cell size by balancing cell elongation and division timing in response to environmental conditions, which is crucial for the optimal use of resources [1]. Changes in cell size are subject to cell functionality optimization [2–5] as well as limitations imposed by the differing kinetics of chemical fluxes across growth conditions [3, 6–8]. The efficiency of nutrient uptake and waste excretion, for example, is determined by the physical dimensions of a cell, including volume, shape, and surface-to-volume ratio [9]. Furthermore, fluctuations in cell size are a key source of noise in gene expression, potentially decreasing cellular functionality and fitness [10–12]. This physiological efficiency, in turn, affects the cell elongation and replication rates, i.e., the fitness of a population. Therefore, controlling cell size is crucial for bacterial fitness, indicating evolutionary pressure to optimize its regulation across nutrient conditions [13].

Understanding cell size under variable nutrient conditions is of particular relevance, as natural growth conditions are rarely constant and require bacterial cells to adjust their size to these changing conditions [14]. With population growth, nutrients are depleted, leading to a slowing and eventually halting of cell growth. Recent developments in cell trapping, tracking and imaging techniques have revealed significant changes in bacterial cell volume during different stages of the growth curve [15–17] with larger cells found during exponential growth (in nutrient-rich conditions) and smaller cells during nutrient-poorer phases [18]. In such time-varying nutrient environments, cell volume appears to be determined by the balance between prioritizing investment in biomass production or cell division [15, 19]. Investigating the dynamics and mechanisms of this balance is important to understand the synchronization of processes connected to growth and division, such as cell elongation, chromosome replication, and protein synthesis [15, 19], which are responsible for the rapid adaptation of bacterial cells to new nutrient environments [14, 19].

In this study, we used single-cell microscopy and modelling approaches to follow cell volume dynamics along the growth curve under various nutrient conditions for two gram-negative rod-shaped bacterial species – *Escherichia coli* and *Salmonella enterica*. We expand the scope of previous investigations by analyzing cell volume distributions over a wider set of nutrient conditions, revealing complex trends in volume dynamics that are not directly related to growth rate but the nutrient environment.

We used microscopy images of rod-shaped bacterial cells at different stages of the growth curve to estimate their volume using recent cell segmentation techniques. We found that cell volume peaked in the early exponential phase and that the volume distribution at this peak could be sustained for several hours by culture dilution with fresh media. In nutrient-limited growth environments, however, cell volume peaks were transient. We found that the timing of the cell volume peaks was not correlated with maximum growth rate but rather the timing of growth rate decrease, i.e., nutrient depletion. Similarly, the magnitude of the peak was affected by nutrient composition. Further, cell volume converged in stationary phase across all conditions, aligning with the hypothesis of a minimum bacterial volume [20, 21]. We propose a heuristic mathematical model to explore the synchronization of growth and division processes necessary to produce and understand the observed cell volume dynamics [22].

## Results

### Cell volume peaks early in the growth curve

First, we studied the growth curve in Lysogeny broth (LB), a standard, but undefined rich medium, by taking consecutive samples for microscopy imaging from an *E. coli* MG1655 culture that was grown under shaking conditions at 37°C (Materials and methods). The first sample was taken from an overnight culture, which had depleted most nutrients and reached stationary phase. These cells were then diluted into fresh LB medium 1:1000 (*t* = 0) and regularly sampled (Fig. 1A-C). Cell volume was estimated from segmented bacterial contours in microscopy images, using the approximation that *E. coli* cells are rod-shaped. We also recorded the optical density of the population (*OD*_600_) as a proxy for biomass at each sampling point to characterize the growth curve dynamics (Fig. 1A) and link them to the estimated mean cell volume (Fig. 1B).

**Figure 1:**
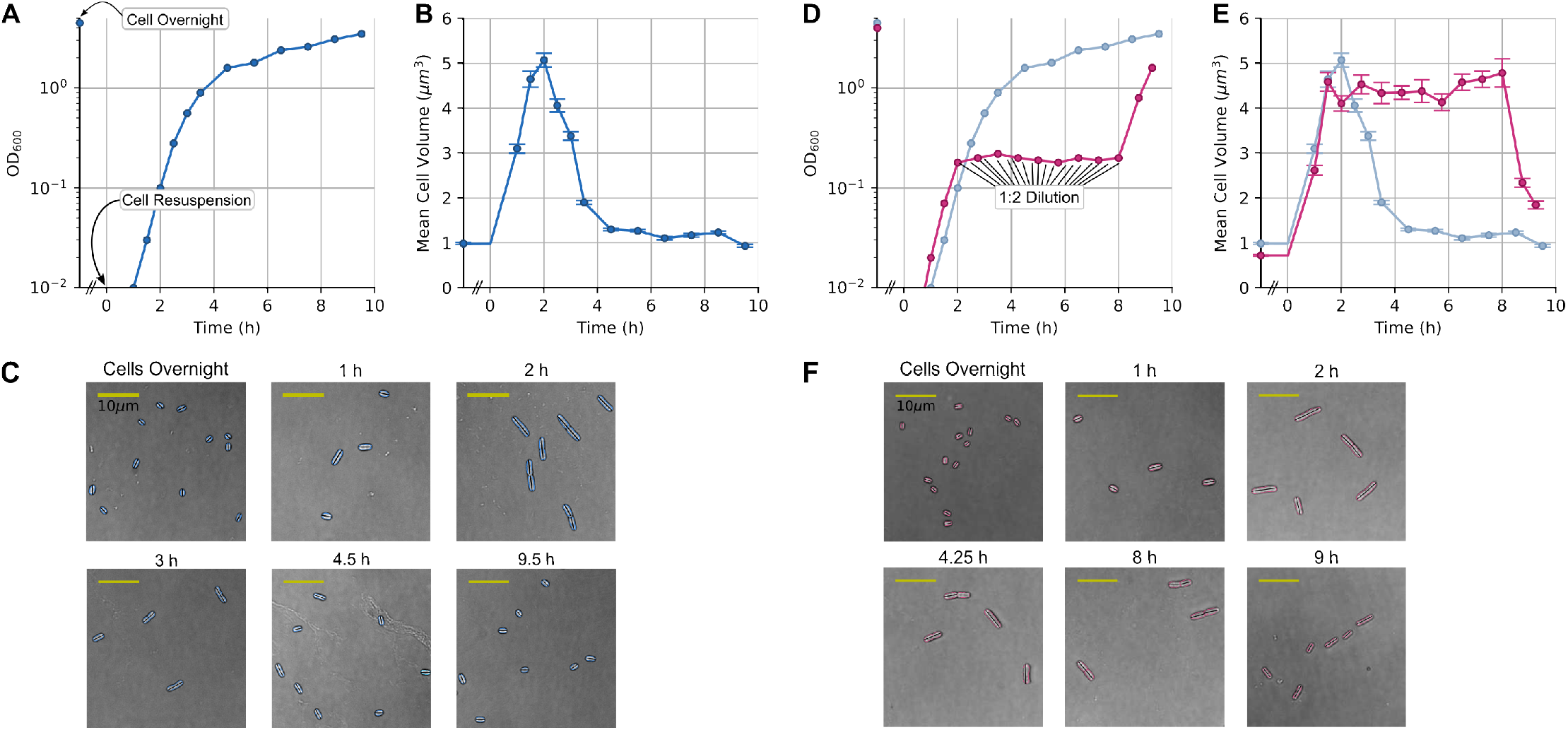
Cell volume dynamics of *E. coli* cells along the growth curve and for sustained exponential growth in rich media. (**A**) Optical density (*OD*_600_) and (**B**) mean cell volume of an *E. coli* culture along the growth curve (blue). (**C**) Examples of cells at different sampling points along the growth curve. (**D**) *OD*_600_ and (**E**) mean cell volume of an *E. coli* culture in sustained exponential growth conditions (pink), which was achieved through consecutive dilutions. The comparison with the regular growth curve is shown in light blue. (**F**) Examples of cells at different sampling points for sustained exponential growth. Colored contours show the cell segmentation and black lines show the dimension used as cell length.

Stationary overnight bacteria had a very small cell volume of ≈ 1*µm*^3^, which increased rapidly to ≈ 5*µm*^3^ over 2h (Fig. 1B,C). This peak volume was maintained only for ≈ 30min and then decreased over the next 2h until it stabilized at a volume slightly larger than that of overnight cells. As previously observed [15], this peak in cell volume results in an opposing, albeit less pronounced, dip in the surface-to-volume ratio (Fig. S1). Analyzing the contributions of cell lengths and widths to volume changes, we observed an ≈ 1.5-fold increase in cell width and an ≈ 2.5-fold increase in cell length. The width increased ≈ 30min earlier than the length, while both decreased over similar time scales (Fig. S1A, S3). Cell length also showed a distinct broadening of the distributions and an increase in noise between 1 and 3h after *t* = 0 (Fig. S2A, S3), coinciding with the time frame of cell volume peak dynamics (Fig. 1B). Notably, the cell culture was still growing for several hours after the cell volume started to decrease. We hypothesized that the observed dynamics can be explained in two ways: either as a transient reaction to cell resuspension and awakening or as a consequence of changes in nutrient levels.

### Sustained exponential growth preserves peak cell volume distributions

To test whether the decrease in cell volume was a transient event after leaving the lag phase or rather dependent on the changing growth environment, we kept the cell culture under sustained exponential growth conditions for 6h. To do so, we diluted the culture 1:2 into fresh medium every 20min, beginning 2h after *t* = 0, i.e., around the time when cell volume peaked (Fig. 1D) (Materials and methods). Under sustained exponential growth, cells maintained a larger mean volume of ≈ 5*µm*^3^ (Fig. 1E) and a broadened volume distribution (Fig. S4) for the entire 6h. When we stopped diluting the culture, the cell volume immediately began to decrease. Cell size homeostasis under steady conditions has previously been reported, mainly from microfluidic experiments [23] and theoretically predicted [24, 25]. Our experiments demonstrate that the transient peak in cell volume in nutrient-limited growth is a physiological adaptation to changes in the growth environment – most likely nutrient depletion – and not a transient effect of growth initiation. This is supported by the fact that cell volume distributions remain broadened as long as the nutrient environment is maintained.

### Poor nutrient conditions lead to almost constant cell volume

To understand how the dynamics of cell volume correlate with population growth, we next investigated cell volume along the growth curve under poor media conditions. We used a defined, minimal medium (M9) with glucose as the sole carbon source [26], which leads to slow growth, as cells need to synthesize amino acids rather than import them from the environment. Under these conditions, cell volume remained roughly constant throughout the growth curve around its stationary phase value of approximately 1*µm*^3^ (Fig. 2A). Cell length increased slightly during growth in M9 glucose while cell width remained constant, overall keeping the cell volume (as well as surface-to-volume ratio) constant (Fig. S1B).

**Figure 2:**
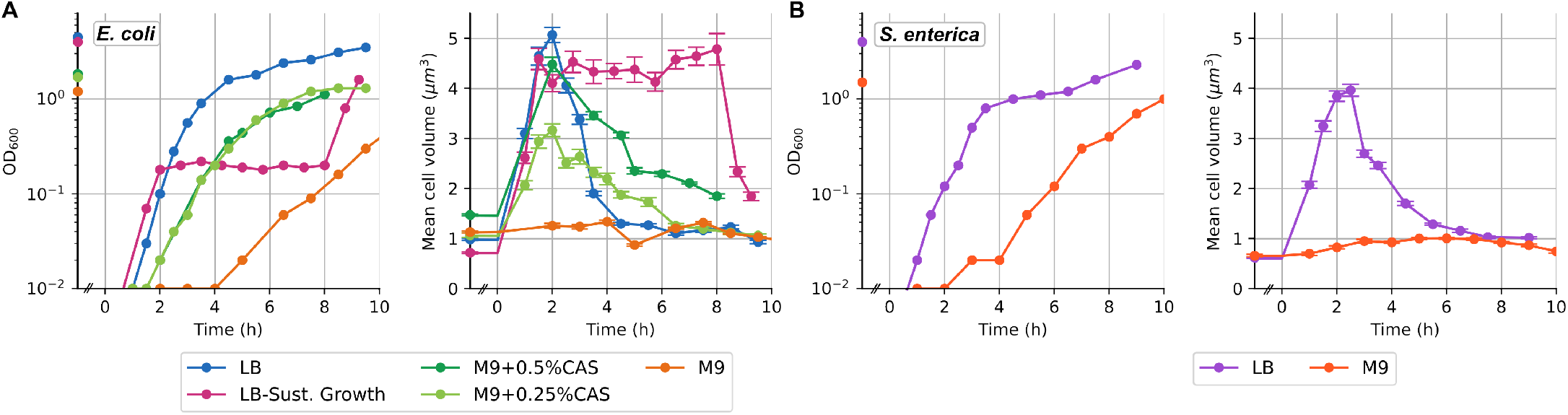
Cell volume dynamics across different nutrient conditions for *E. coli* and *S. enterica*. *OD*_600_ of the cell culture (left) and mean cell volume (right) over time in different conditions: (**A**) rich to poor nutrient growth curves (blue, dark and light green, orange) and sustained exponential growth (pink) for *E. coli*; (**B**) rich (purple) and poor (red) nutrient growth curves for *S. enterica*.

As cell volume dynamics in rich and poor nutrient media differed drastically, we repeated these experiments in another gram-negative microorganism, *Salmonella enterica* LT2. As *S. enterica* is also rod-shaped and divides by symmetric binary fission, we would expect similar cell volume regulation as in *E. coli*, even though this has not been extensively studied before. Indeed, we found peak dynamics in rich (LB) media and almost constant cell volume in minimal (M9 glucose) media (Fig. 2B, S1A) as seen with *E. coli* (Fig. 2A, S1A).

### Peak cell volume is dependent on nutrient composition

The contrasting cell volume dynamics in rich and poor nutrient conditions led us to investigate the nutrient dependence of cell volume changes. We studied *E. coli* in two intermediate nutrient conditions: defined M9 medium with glycerol, supplemented with 0.25% or 0.5% casamino acids (partially digested amino acids). Surprisingly, both media resulted in similar growth curves but different cell volumes (Fig. 2A). The timing of the cell volume peak in both supplemented M9 media was very similar (around 2h after resuspension), but the mean peak volume was much larger for 0.5% casamino acid supplementation compared to 0.25%. Cells grown in M9 with 0.5% casamino acids had a similar peak cell volume to that seen with LB, while growth in 0.25% casamino acid supplementation resulted in a peak volume approximately halfway between poor and rich media (≈3*µm*^3^) (Fig. 2A). Similarly, when calculating the surface-to-volume ratio, we found a much smaller change for M9 supplemented with 0.25% casamino acids, although the difference was not as pronounced as for cell volume (Fig. S1B). Generally, for LB and M9 supplemented with casamino acids, the surface-to-volume ratio decreased during the first 2h of growth and then increased during the transition to stationary phase, opposite to cell volume changes, but slower and with a lower magnitude of change (Fig. S1). The surface-to-volume ratio in M9 glucose remained relatively constant throughout the growth curve and was higher than in all richer media. This is correlated with the cell width in M9 glucose being consistently smaller than in the other media (Fig. S1). Overall, we found a strong nutrient dependence of the maximum cell volume and the cell width, whereas the timing of the volume peak was similar.

### The ratio between biomass (*OD*_600_) and cell numbers (CFU) shows similar peak dynamics as cell volume

As we found a highly dynamic cell volume behaviour over the exponential growth phase in most media, we realized that this could affect our measurements of population growth. Population growth or increase in biomass of bacterial cultures in liquid media is commonly measured through optical density (*OD*_600_) [28], assuming that *OD*_600_ correlates linearly with biomass and that biomass correlates approximately linearly with cell density. However, this relationship is highly dependent on the assumption that the biomass contributed by each cell is con-sistent over time [29]. The drastic changes in cell volume observed within the first hours of growth indicate that this assumption does not hold under most media conditions.

To test the dynamic relationship between cell numbers and biomass, we sampled *E. coli* cultures grown in LB over 4h after resuspension of overnight cultures and determined colony forming units (CFUs) as well as biomass (*OD*_600_) (Fig. 3A,B). We used three biological replicates plus four technical replicates each for CFU counts, taking measurements every 30min (Materials and methods). We fitted curves through these measurements and calculated the biomass per cell as *OD*_600_/CFU using inference based on Gaussian processes [27]. We found a similar peak pattern as for cell volume, although the peak occurs slightly earlier, at 1.5h instead of 2h after resuspension (Fig. 3C). The peak pattern agrees with the idea that our estimate of biomass per cell is approximately proportional to the average cell volume in the culture at any given time point.

**Figure 3:**
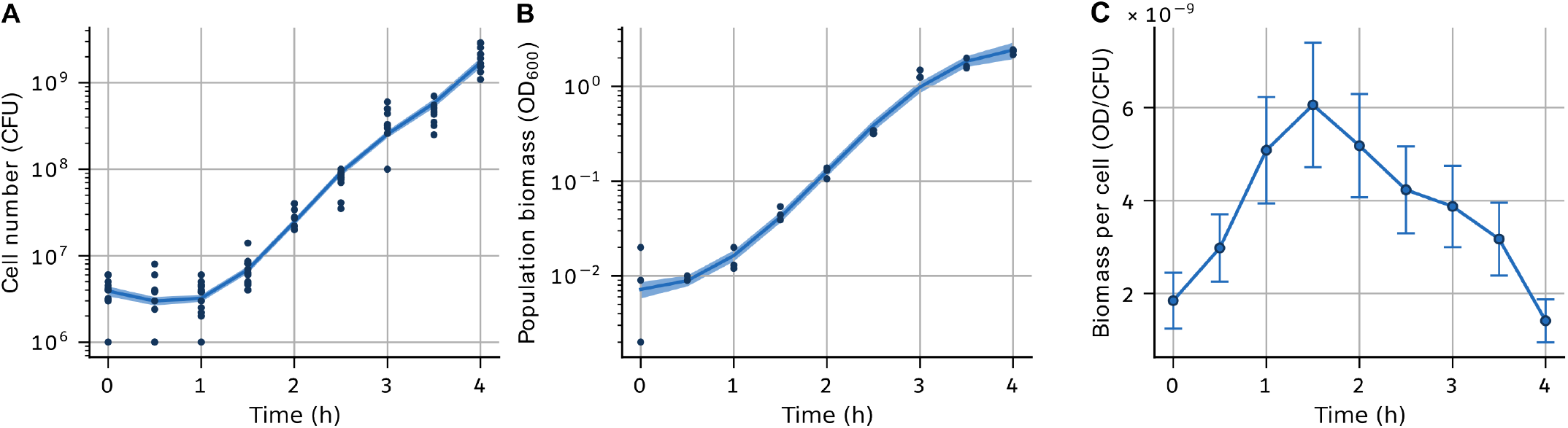
The ratio between cell numbers and biomass shows a similar pattern as cell volume over the growth curve. (**A**) Cell numbers as colony-forming units (CFU) (three biological replicates with four technical replicates each) and (**B**) biomass as *OD*_600_ (three biological replicates) along the growth curve in LB, where lines show the best-fit curve to measurements. (**C**) The ratio between biomass (*OD*_600_) and cell numbers (CFU) over the growth curve. Confidence intervals were estimated using inference methods based on Gaussian processes [27].

The peak in the ratio between *OD*_600_ and CFU is caused by a discrepancy between cell and population growth. Although cell elongation began soon after resuspension as visible in the increase in *OD*_600_ after 0.5-1h, cells were not yet dividing, meaning that CFU counts only increased after 1.5-2h. Hence, biomass increased soon after inoculation, but population growth was delayed by ≈ 1h under these conditions. Conversely, after 2.5h, CFU numbers still indicate exponential population growth, while biomass growth is already levelling off, indicating a decrease in biomass per cell, i.e., cell volume (Fig. 3).

### A heuristic model relates changes in cell size to dynamic resource allocation

The relationship between population growth (cell division) and biomass growth (cell growth) is dynamic, regulating cell volume under varying nutrient conditions. To understand these changes in cell division and growth over the growth curve, we used a simple discrete-event model. As the model does not take cell shape into account, we are going to refer to cell size instead of cell volume here, but for our results, the two terms can be seen as equivalent. We describe the change in cell size *s*(*t*) by assuming that cells grow exponentially at a time-dependent rate *µ*(*t*):

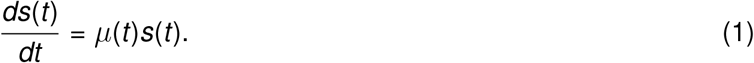

We can estimate *µ*(*t*) from the increase in biomass in our experiments (Fig. 4A,B): we fitted a logistic curve to the *OD*_600_ measurements and used it to calculate the growth rate over time (Materials and methods). As expected, *µ*(*t*) is high after resuspension (*t* = 0) and then decreases over the growth curve until it approaches zero with entry into the stationary phase. For simplicity, we ignore the slow growth phase during the transition into the stationary phase and focus on the main trends along the curve. Strikingly, *µ*(*t*) is well described by a constant growth rate until ≈ 2h after cell resuspension in all rich-nutrient conditions, which is the time point after which volume typically decreased as well (Fig. 4B).

**Figure 4:**
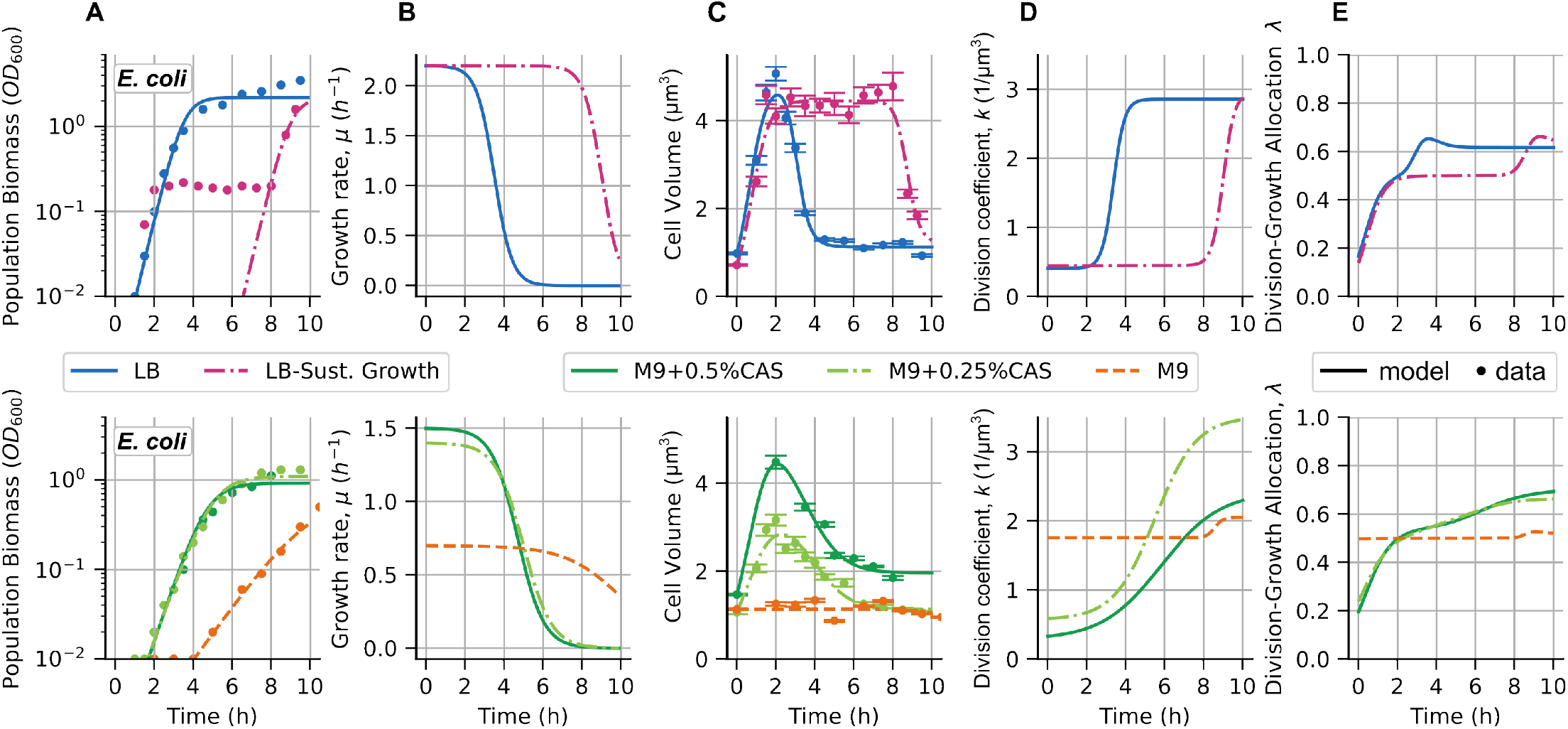
A model of cell volume dynamics shows the modulation of division rate and resource allocation over the growth curve. (**A**) Biomass (*OD*_600_) over time for various nutrient conditions as shown in different colours; points show measurements (Fig. 2A), solid lines are the fit to a logistic curve. The biomass growth for sustained exponential growth conditions (pink line) was approximated using the biomass growth in LB. (**B**) Growth rates obtained from fitted curves. (**C**) Comparison of the mean cell volumes shown in Fig. 2A (points and error bars) and the prediction of the mean-field model of cell volume regulation (solid lines). (**D**) Dynamics of the division rate *k* over the growth curve in different nutrient conditions. (**E**) Dynamics of the division-growth allocation parameter *λ* over the growth curve in different nutrient conditions.

Cell division can be described by halving the cell size at discrete time points, as determined by the division rate. We assume that the division rate depends on cell size, resulting in *adder* size control if the division rate is directly proportional to cell size [30, 31]. *Adder* size control means that, on average, cells add a constant size between divisions, and is the cell size control model generally associated with *E. coli* [32–34] and other bacterial species [35–38]. Hence, we assume that the division rate scales with the cell size as well as the growth rate following *k* (*t*)*µ*(*t*)*s*(*t*), where *µ*(*t*)*s*(*t*) is the rate of change in cell size and *k* (*t*) is defined as the time-dependent division coefficient. The division event can then be described as a discrete jump:

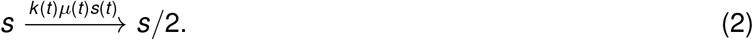

Using the mean-field approach (Materials and methods), we can now describe the dynamics of the mean cell size ⟨*s*(*t*)⟩ (⟨.⟩ indicating the mean at any given time point) through growth and division:

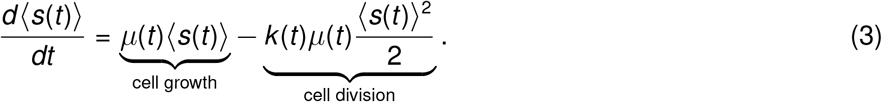

For constant *k*, cell division and growth are synchronized and the mean cell volume will approach the steady-state value: 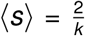. In this case, the mean cell volume is independent of the growth rate as the effect of *µ* on growth and division balances out.

However, our experiments showed that cell volume, and therefore the division coefficient *k*, was not constant over the growth curve under most nutrient conditions (Fig. 4B,C). We approximated the dynamics of *k* using a sigmoidal function over time (Eq. (14)) and solved the equation for mean cell size (Eq. (3)) numerically by using our measurements for cell size (volume) and cell growth (biomass) dynamics over the growth curve (Materials and methods). We found that the division coefficient *k* starts low, increases when the growth rate begins to decrease, and then plateaus at its highest value when the growth rate also plateaus (Fig. 4B, 4D). The steepness of *k* increase differs between nutrient conditions and highlights that a combination of growth rate and cell volume determines division (see, for example, M9 supplemented with 0.25% and 0.5% casamino acids in Fig. 4B-D).

How then does the relationship between cell growth and division determine cell volume? Changing nutrient conditions require bacterial cells to dynamically navigate the balance between cell growth and division by adjusting their proteome allocation between biomass accumulation and division machinery synthesis [19]. Here, we avoid the complexities of modelling proteome allocation change and focus on understanding the qualitative changes in division-growth allocation. We can calculate the division-growth allocation coefficient *λ* as the proportion of division compared to all size regulation processes (i.e., division and growth) (Materials and methods):

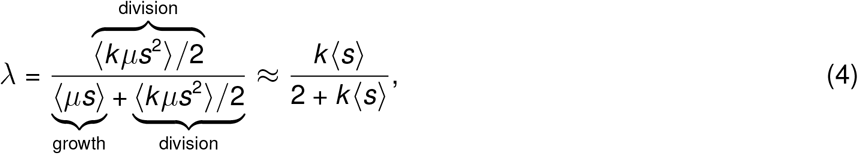

*λ* describes the proportional allocation of resources to division or growth, with all resources allocated to division for *λ* = 1 and all resources allocated to growth for *λ* = 0. The allocation is exactly 50:50 for *λ* = 0.5, which we can see for sustained growth in LB as well as for M9 with glucose, where the division coefficient and cell size remained constant (Fig. 4E). While *λ* starts to increase again in the sustained growth experiment after the dilutions are stopped, for M9 with glucose, the almost perfect balance of division and growth led to constant cell volume over 10h of growth. In all other nutrient conditions, division-growth allocation started from ≈ 20% (balanced towards growth) and increased to 60-70% (balanced towards division) when entering stationary phase (Fig. 4E), which is qualitatively consistent with studies using more sophisticated proteome allocation models [19]. Interestingly, growth rates and division-growth allocation are very similar for M9 supplemented with 0.25% or 0.5% casamino acids, but the division coefficient *k* has to differ to result in the experimentally observed differences in cell volume. This means that slight differences in nutrients can change the relationship of growth rate and cell volume with the propensity for division.

## Discussion

Our study reveals general trends of cell volume dynamics and its link to growth rate and nutrient conditions over the growth curve. We found that the timing of cell volume dynamics was not directly correlated with maximum growth rate: in all nutrient-rich media (LB, M9 supplemented with casamino acids) cell volume peaked ≈ 2h after cell resuspension. The magnitude of the cell volume peak correlated however with the nutrient composition, with richer media leading to larger maximum cell volumes, which was most clearly visible across different M9 conditions (Fig. 2). While richer media also led to higher growth rates, growth rate and cell volume seemed to be related through different functions with the nutrient composition of the growth medium (Fig. 4B,C).

In nutrient-rich media, growth rate estimates showed a decrease after 2-3h. Across nutrient conditions, there is a coincidence between the instant when this growth started to decrease and the peak in cell volume (Fig. 4). This would imply that a decrease in growth rate, likely caused by the onset of nutrient depletion, determines the cell volume peak. The rate of volume decrease toward the common cell volume in stationary phase was different between nutrient conditions, but not directly correlated with growth rate or nutrient composition. Generally, the increase and decrease in cell volume were steepest in LB, where we also saw the fastest growth rates. The trends in cell size dynamics were less clear for M9 supplemented with 0.25% or 0.5% casamino acids, where growth rates were very similar along the growth curve, but the maximum cell volume and cell volume dynamics were markedly different (Fig. 2A). This surprising independence of cell volume changes and growth rate can be understood through our model: growth rate can affect both processes involved in cell volume regulation, growth and division, differently, potentially as a function of the nutrient conditions. In our model, size regulation through division is quantified by the division coefficient *k* (Eq. (3)), which can be understood as how nutrients (and potentially other physiological factors) are translated into the division rate.

Cell volume is just one of the dimensions that we can consider for cell size dynamics. Other measures are cell length and width, which can be determined from the segmentation masks of the microscopy images [39]. We found that cell width increased earlier than cell length, with a delay of ≈ 30min (Fig. S1), which is consistent with previous studies [15, 40]. Surprisingly, there seemed to be more variation between nutrient conditions for cell width changes, although changes over the growth curve were greater in cell length (Fig. S1). The difference in cell volume dynamics between cells grown in M9 supplemented with 0.25% and 0.5% casamino acids, for example, was mainly because cells were significantly wider in the latter, while lengths were similar (Fig. S1). Such changes are particularly interesting, as bacteria can use changes in cell shape, specifically changes in surface-to-volume ratio, as a means of antibiotic resistance by reducing the intracellular or membrane concentration of an antibiotic [41]. Hence, understanding changes in cell length and width across nutrient conditions is important to determine the potential for cell-shape-based antibiotic resistance.

Poor medium (M9 with glucose) presented a special case, in which cells did not show appreciable changes in cell volume or surface-to-volume ratio over the growth curve (Fig. S1B,C). This was mainly due to the constant cell width throughout the growth curve, although the cell length increased slightly (Fig. S1B,C). Interestingly, cells were thinner and longer than cells with similar volume in other growth conditions (which usually occurs during their transition to stationary phase) (Fig. S1B,C). In *S. enterica*, the increase in cell length in M9 with glucose was more pronounced than in *E. coli*, giving an overall slight increase in cell volume, which only decreased again ≈ 7h into the experiment (Fig. S1) at which point the growth rate also started to decrease (Fig. 4B). These results suggest that the conserved 2h peak in all other nutrient conditions was coincidental and that the peak is related to the change in growth rate, potentially caused by nutrient depletion. However, given the small change, lack of a proper peak in volume dynamics and very slow growth rate, the poor medium does not seem to allow for conclusive evidence about the regulation of cell volume dynamics. Furthermore, our model showed an almost perfect 50:50 balance of resource allocation between growth and division over most of the growth curve in M9 with glucose (Fig. 4E), suggesting little change in size regulation.

While cells in minimal media already started with a 50:50 balance of cell growth and division allocation (*λ*), this allocation usually started from a low value under other nutrient conditions. In these conditions, cells showed a growth-biased allocation that rose toward 50:50 allocation within the first 2h. This dynamics is indicative of drastic changes in cell volume during the first 2h of the growth curve, but not necessarily of cell division. In fact, our results showed that the ratio between cell number (CFU) and biomass (*OD*_600_) is non-monotonic (Fig. 3). This means that during the first ≈ 1.5h, the increase in biomass appeared to be mainly due to an increase in cell volume in individual cells, but not an increase in the number of cells. From these measurements, we estimate that the maximum biomass per cell was reached after ≈ 1 doubling time. The peak in cell size occurs half an hour later, which would amount to 2-3 cell doublings. The difference in those estimates could be due to inaccuracies in OD measurements and difficulties in determining when cells have finished dividing from microscopy images. Overall, the ratio between OD and CFU showed a peak similar to that of cell volume, which makes sense since this ratio can be associated with the biomass per cell, i.e., is roughly equivalent to cell volume (Fig. 1A, 3C). This highly non-linear relationship can lead to significant overestimation of cell numbers based on optical density or turbidity, yet these measures are commonly used in microbiology experiments. Non-linearity also makes the accuracy of cell number estimates using *OD*_600_ highly dependent on the use of the same conditions and the timing of the measurements to make them comparable, which could be alleviated by using calibration curves to establish the *OD*_600_-CFU relationship for each nutrient condition [42, 43].

The main part of our analysis focused on the mean cell size behaviour, but we also observed interesting patterns in the random size variability between individual cells (Fig. S2). We observed that the awakening of cells after resuspension led to a broadening of the cell volume distribution that lasted throughout most of the peak dynamics (Fig. S3). This variation appeared to come from variation in cell length rather than cell width (Fig. S2, S3), indicating heterogeneity in the timing of the first division between cells. Surprisingly, the increase in cell volume and length variation did not seem to be a transient phenomenon, as sustained exponential growth conditions preserved not only the maximum cell volume but also the broadness of the distribution (Fig. S4). Using a stochastic model of cell size regulation, we found that sustained high variation in cell volume during sustained exponential growth (Fig. S5) can be explained if the noise in size regulation has a weak dependence on cell size itself. In general, the increase and decrease in the mean and variation of cell volume appear to be related to a change in the nutrient environment.

Our study opens intriguing avenues for further experiments and modelling. First, the convergence to a common cell size during stationary phase raises important questions. Current models based on growth-division allocation capture general trends in cell size dynamics but fail to explain the robustness of this convergence across growth conditions [19]. Possibly, the common cell size corresponds to a minimal threshold size below which cell division is disabled [21, 44]. Since this minimum cell size is related to the initiation of chromosome replication, cell size dynamics in fluctuating environments could be examined by labelling the replication origins [20]. Second, the potential size dependence of noise is intriguing and a deeper understanding of its origin could shed light on the constraints governing cell size regulation. Previous studies indicate that understanding stochasticity in cell division mechanisms allows us to distinguish between various model types that predict similar mean dynamics [44]. Coupling such modelling frameworks with time-dependent dynamics has proven useful for characterizing cell size homeostasis in diverse systems such as cancer cells [45], yeast [46, 47] and different bacterial species [25]. Third, modelling the growth curve itself is challenging. While we used a logistic model to describe the population dynamics, models such as the Gompertz model provide a more accurate fit in certain contexts (e.g., tumour growth) [48]. Gompertz models are particularly relevant when accounting for multiple cell phenotypes, such as quiescent and proliferating subpopulations, which vary in proportion along the growth curve [49], or subpopulations metabolizing different nutrient sources, arising from diauxic growth [50]. Including these subpopulations could enhance the predictive power and biological relevance of our approach. Our work also highlights the importance of considering how nutrient shifts affect resource allocation for predicting cell volume regulation across changing environments [19], similar to how these shifts influence growth transition kinetics [6]. Mechanistic models of nutrient consumption could be coupled with experiments that modulate the type and concentration of nutrients (e.g., nitrogen and carbon sources) to gain a more comprehensive picture of cell volume mean and noise dynamics [51]. Further, RNAseq at multiple time points (e.g., 1 and 2 hours after resuspension) could link gene expression patterns with cell volume changes, offering deeper insights into cellular responses to nutrient shifts [52, 53]. These approaches could serve as a starting point for exploring size-dependent noise and population heterogeneity, addressing the questions highlighted above.

In this study, we investigated cell volume dynamics in two rod-shaped bacteria, *E. coli* and *S. enterica*, along the growth curve, revealing a strong dependence of cell size regulation on nutrient conditions. Our findings reveal the remarkable plasticity and adaptability of bacterial cells in regulating their size in response to environmental changes by balancing resource allocation between cell growth and division in response to nutrient availability and potentially nutrient type. The relationship between cell size and nutrient conditions is highly relevant to understanding the behaviour of natural microbial communities, but also clinical phenotypes, such as resistance to antibiotics and persister cells [17, 41].

## Materials and methods

### Bacterial growth conditions and media

Experiments were carried out using the *E. coli* K-12 substrain MG1655 and the *S. typhimurium* LT2 derivative, TH437. Bacterial cells were grown overnight at 37C with shaking and aeration in 15 ml culture tubes filled with 2 ml of growth medium. Minimal media consisted of 1X M9 salts, 1mM thiamine hydrochloride, 0.4% glucose, 0.5% or 0.25% casamino acids, 2mM MgSO4, 0.1mM CaCl2. M9 salts, casamino acids and LB media were autoclaved, and other ingredients were sterilized with filter sterilization. Overnight cultures were diluted 1:1000 v/v into 10ml of the same medium and grown on the shaking incubator with aeration at 37C. Samples were taken at 30- or 60-min intervals and imaged under the microscope. At each sampling point, *OD*_600_ measurements were taken with a spectrophotometer using either 1ml of the undiluted culture or at later time points, by diluting the culture in the same medium to an *OD*_600_ less than 1 to obtain accurate measurements.

For cultures that were kept in an early exponential phase in rich media, half of the culture was diluted with the same amount of fresh media every 20min and sampled for imaging at every second dilution.

### Single-cell microscopy

Between 1 and 6 *µl* of the sample (diluted 2- or 4-fold at later time points to avoid cell clusters) were spotted on an agar pad (made of minimal media with 1.5% agarose) and left to dry for a few minutes. The agar pad was inverted and placed into a microscopy dish (*µ*-Dish 35mm, low; Ibidi). The microscope dish was mounted on an inverted microscope stage (Eclipse Ti2-E, Nikon) and bright-field images were taken with a Nikon DS-Qi2 camera using a 100× oil immersion objective (Plan Apo *λ*, N.A. 1.45, Nikon).

### CFU count plating

For cell number counts, we diluted overnight cultures of *E. coli* K-12 substrain MG1655 1:1000 into fresh LB media. 3 biological replicates were measured every 30min for 4h to obtain *OD*_600_ via a spectrophotometer (see above) and CFU by plating. For plating, we used the running droplet method where 10ul of each sample dilution (10^0^ − 10^−7^) are dropped onto a square plate, which is then tilted so that droplets can run down to half of the plate. We plated each dilution twice and used all dilutions with countable colony numbers for analysis.

### Segmentation and analysis of microscopy data

We estimated the cell volume from the bright-field microscopy images by segmenting cells using the pixel-classification module of Ilastik (v.1.3.3) [54]. As each nutrient medium influenced the cell shape differently along the growth curve, we trained an Ilastik neural network for each condition. The training process involved outlining an arbitrary set of cells to define the contours and conducting manual checks to resolve instances where cell clusters posed segmentation challenges. We devoted special attention to exponential phase images, to differentiate between cells that had not yet undergone division and cells that had undergone division but remained physically connected. This is exemplified in Figure 1C and 1F at the 2-hour mark.

From the estimated contours we measured the projected area *A*_*p*_, which corresponded to the number of pixels inside the contour times the pixel area (≈ 0.005*µm*^2^*/pixel*). We defined the cell length *L* as the longest side of the minimum-bounding rectangle of the contour. The projected area *A*_*p*_ and the cell length *L* are related to the effective cell width *w* through the projected area of a capsule:

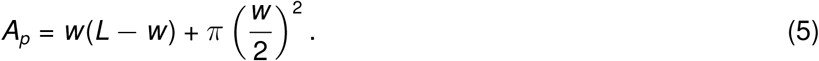

Thus, from *A*_*p*_ and *L, w* is estimated by solving Eq. (5). The cell surface area *A* and volume *V* can then be estimated from *L* and *w* as follows:

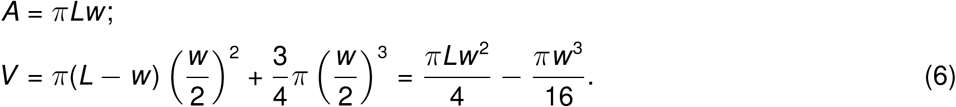

After obtaining these dimensions for all segmented cells, we filtered the outliers (usually imaging artefacts) considering only cells satisfying the following criteria:

- Cell width greater than 0.35 *µ*m.
- Cell length greater than 1.05 *µ*m and less than 10 *µ*m.
- Cell aspect ratio (*L/w*) greater than 1 and less than 7.
- Cell area greater than 0.73 *µm*^2^.

Further, for each time point, we applied a final outlier filtering step. We discarded any cells where the deviation in the log-transformed cell volume from the log-transformed population mean cell volume exceeded three times the log-transformed population standard deviation of the cell volume. This assumes that the cell size distributions follow a log-normal distribution and therefore, this range should include approximately 99.7% of the data. Statistics and confidence intervals were estimated using Bayesian methods [55].

### Estimation of OD and CFU time curves

To estimate the ratio OD/CFU over 4h after resuspension, we ran an inference algorithm [27] separately on the replicates of cell number and population biomass respectively. As a result, we obtained the most probable trajectory of the mean with its 95% confidence interval (Fig. 3A, B). Given the most probable mean OD *x* with its confidence interval Δ*x*, and the most probable CFU *y* with its confidence interval Δ*y*, the most probable ratio *z* and its confidence interval Δ*z* were estimated using the formulas:

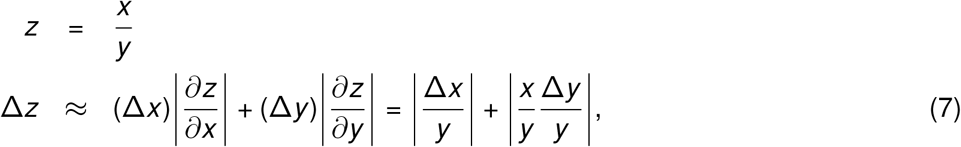

Standard methods for propagation of uncertainty were used for this estimation.

### Mean-Field cell size dynamics

A more detailed approach to the theoretical model was published recently [30]. Briefly, assuming that cells grow following the dynamics in Eq. (1) and halve their size at each division, the expected value of any arbitrary function of cell size *f* (*s*) follows the dynamics:

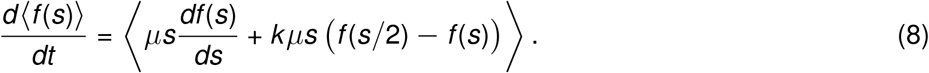

To estimate the *n*-th moment of cell size, we replaced *f* (*s*) = *s*^*n*^ and obtained the differential equation governing the dynamics of ⟨*s*^*n*^⟩:

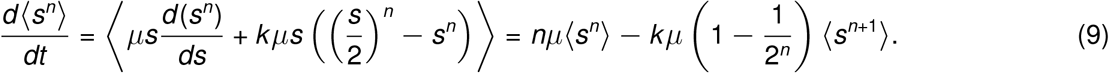

The main issue with equation (9) is that the dynamics of ⟨*s*^*n*^⟩ depend on ⟨*s*^*n*+1^⟩. This problem is known as unclosed moments dynamics [56] and has been explored recently [57]. The mean-field approximation neglects the random fluctuations of *s* and its correlation with other variables. Therefore, we approximate the second-order moment as ⟨*s*^2^⟩ ≈⟨*s*⟩^2^ and ignore the equations for higher-order moments to obtain the equation (3) in the main text.

### Fitting procedure for the division coefficient

To simplify our approach, we first fitted the OD curves to a logistic curve of time. This simplification assumes that the growth rate is proportional to the nutrients and that the rate of nutrient depletion is proportional to the biomass. In summary, we assumed that the biomass *B* follows:

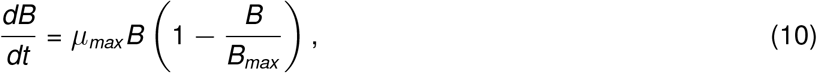

where *µ*_*max*_ and *B*_*max*_ are the maximum growth rate and the carrying capacity of the growth medium, respectively. We adjusted the OD dynamics to the solution of (10):

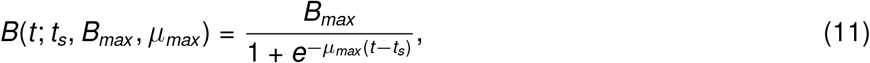

where the free parameter *t*_*s*_, depending on the initial conditions of *B* can be interpreted as the time for entering stationary phase. The parameters *t*_*s*_, *B*_*max*_ and *µ*_*max*_ were fitted to data by minimizing the error function:

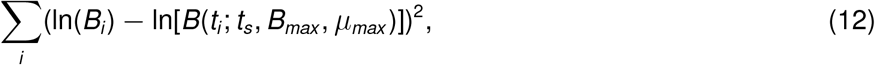

in which *B*(*t*_*i*_ ; *t*_*s*_, *B*_*max*_, *µ*_*max*_) is calculated from the equation (11) at *t* = *t*_*i*_ for each *i* -th point. Note that the error function is the Euclidean distance between the logarithm of the data and the logarithm of the model with *t*_*s*_, *B*_*max*_, *µ*_*max*_ as fitting parameters. After fitting, from the equation for biomass over time (11) we could then obtain the growth rate over time:

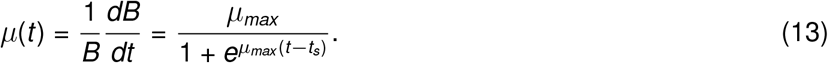

The division dynamics were assumed for simplicity to be described by the division coefficient that follows the dynamics:

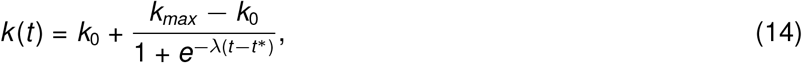

with four free parameters: the basal coefficient *k*_0_, the maximum coefficient *k*_*max*_, the rate of change *λ* and the time of reallocation *t*^*^ is the value of time where *k* (*t*) reaches the halfway point between *k*_0_ and *k*_*max*_. Proposing a set of parameters *k*_0_, *k*_*max*_, *λ* and *t*^*^, we can numerically integrate the mean cell size ⟨*s*⟩ from the differential equation (3) with initial condition ⟨*s*⟩|_*t*=0_ = *s*_1_. The mean cell size at time *t* = 0 was taken as the mean cell size from overnight cultures. The optimized parameters for *k* (*t*) were selected after a comparison between the observed mean sizes {⟨*s*⟩_1_, … ⟨*s*⟩_*N*_} and the predictions {⟨*s*(*t*_*i*_)⟩, …⟨*s*(*t*_*i*_)⟩_*N*_}, where ⟨*s*(*t*)⟩ is the numerical solution of Eq. (3). This optimization minimized the error function:

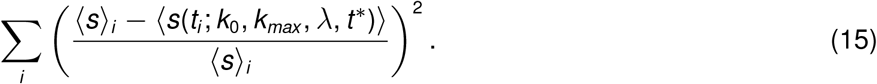

Note that this error function corresponds to the square of the difference in mean cell size between theory and experiment divided by the observed mean cell size.

## Author Contributions

- **Conceptualization:** C.I. C.N., A.S.
- **Investigation:** C.I.
- **Data Curation:** C.N.
- **Formal Analysis:** C.I. C.N., A.S.
- **Funding Acquisition:** C.I., A.S.
- **Methodology:** C.I. C.N., A.S.
- **Resources:** C.I., A.S.
- **Software:** C.N.
- **Supervision:** A.S.
- **Visualization:** C.I., C.N.
- **Writing:** C.I., C.N., A.S.
- **Writing – Review Editing:** C.I., C.N., A.S.

## Acknowledgements

We thank M. Lagator for helpful comments on the manuscript and M. La Fortezza for his support with the microscopy setup. This work was supported by the Wellcome Trust (225565/Z/22/Z) and an ETH Zurich Post-doctoral Fellowship (19-2 FEL-74) received by CI. AS and CN acknowledge the support of NIH-NIGMS through grant R35GM148351.

## Artificial intelligence

The authors declare that they did not use generative AI and AI-assisted technologies in the writing process.

## Data accessibility statement

The data set and data processing scripts are publicly available at https://doi.org/10.5281/zenodo.13208281 [58].

