## Supplementary Material for "Bacterial cell size modulation along the growth curve across nutrient conditions"

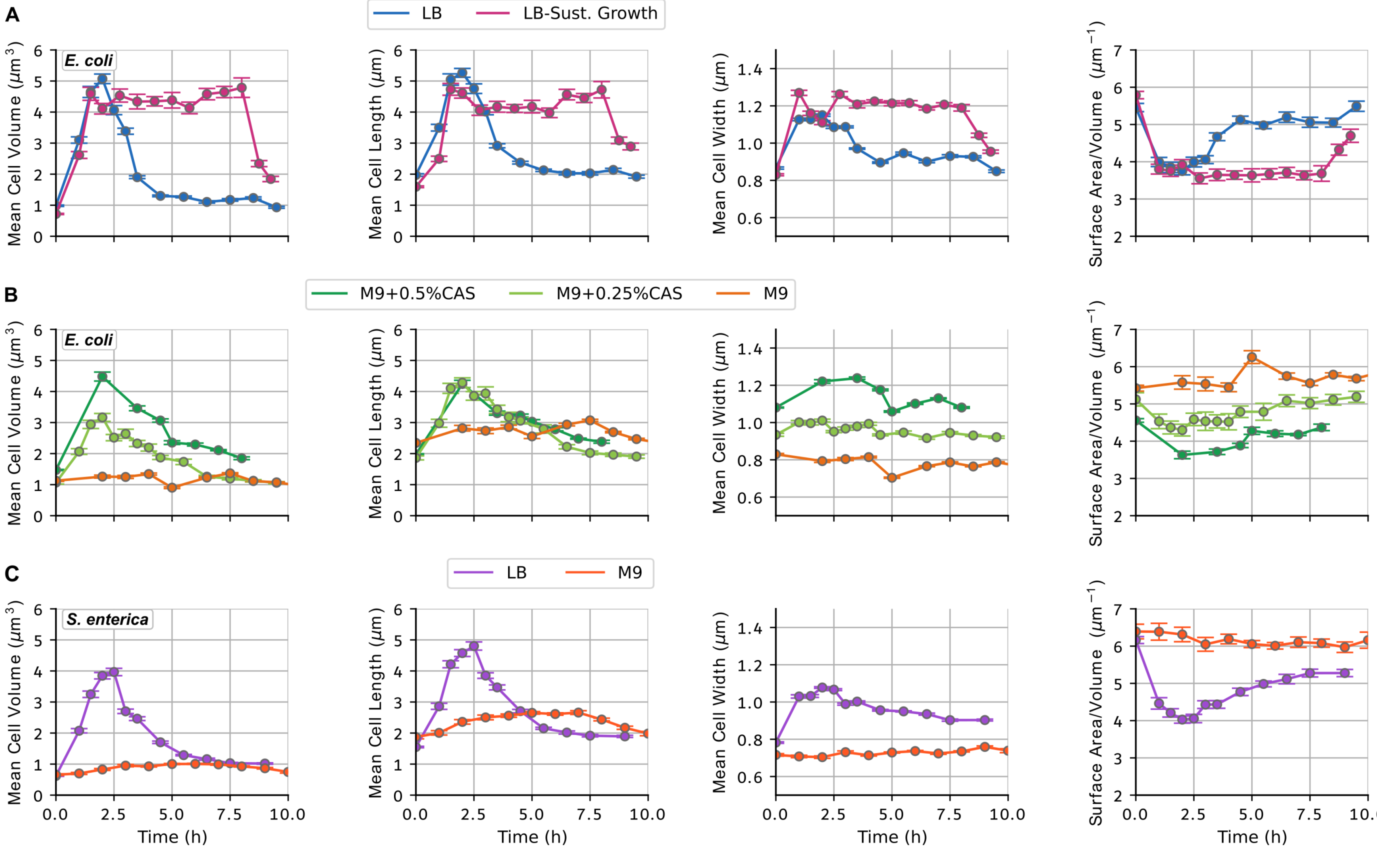


Figure S1: **Cell volume, length, width and surface-to-volume ratio dynamics across different nutrient conditions for *E. coli* and *S. enterica*.** Mean cell volume, length, width, and surface-to-volume ratio of (**A**) *E. coli* cell cultures at different sampling points over time in LB (nutrient-limited in blue and sustained exponential in pink), (**B**) *E. coli* cell cultures at different sampling points over time in M9 glucose (orange) or M9 glycerol supplemented with 0.25% or 0.5% casamino acids (light and dark green), or (**C**) *S. enterica* cell cultures at different sampling points over time along rich (purple) and poor (red) nutrient growth curves. Overall, we found that changes in cell length were more pronounced than changes in cell width. Cell width was consistently smaller in minimal media with glucose than in richer media. Interestingly, for minimal media with different supplements, cell width was more distinct than cell length (B). LB media showed the biggest change in cell length and led to the strongest drop in the surface area-to-volume ratio.

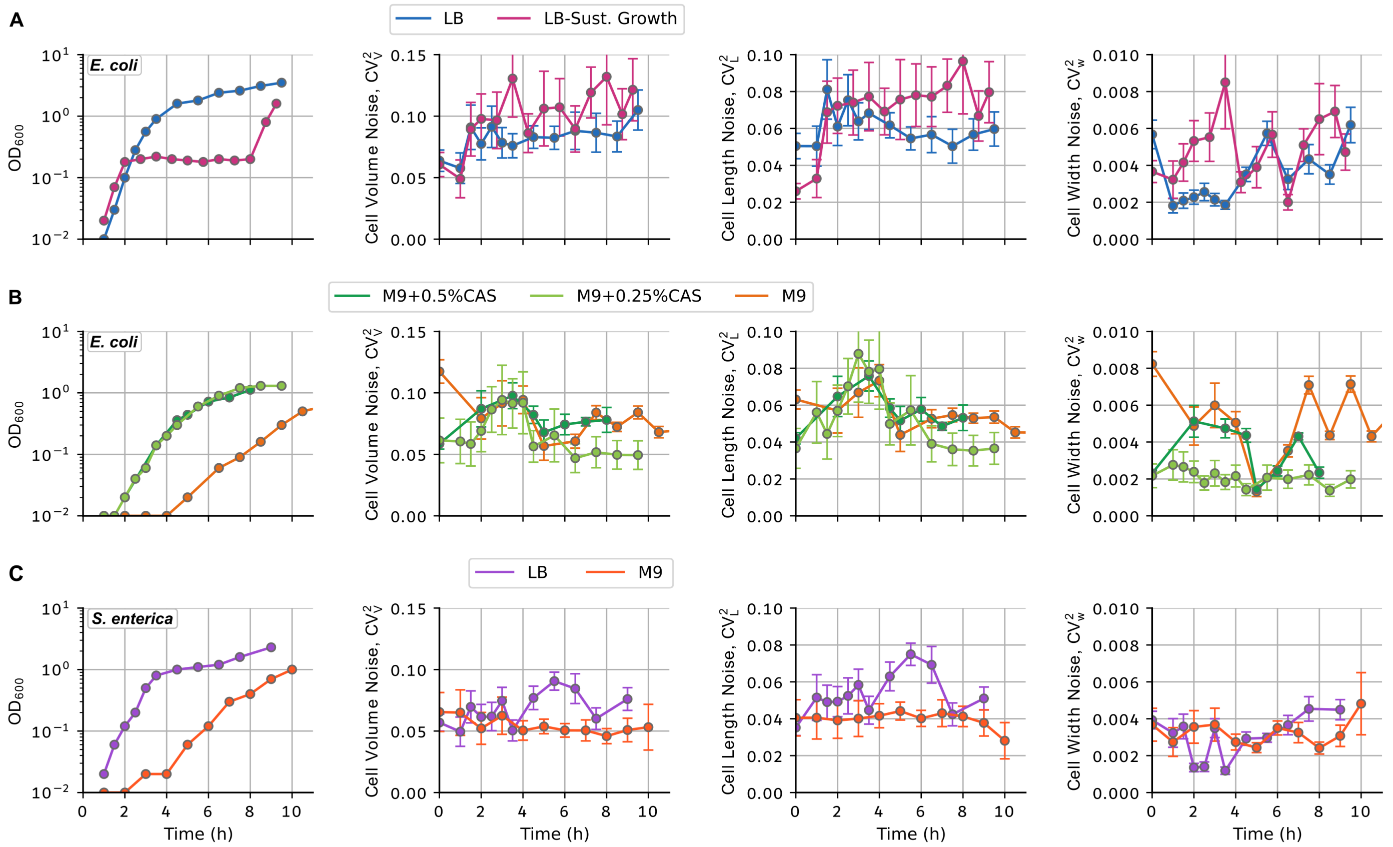


Figure S2: **Noise in cell volume, length and width across different nutrient conditions for *E. coli* and *S. enterica*.** $OD_{600}$ and noise (quantified by the coefficient of variation squared $CV^2$) of cell volume, length and width for (**A**) *E. coli* cell cultures at different sampling points over time in LB (nutrient-limited in blue and sustained exponential in pink), (**B**) *E. coli* cell cultures at different sampling points over time in M9 glucose (orange) or M9 glycerol supplemented with 0.25% or 0.5% casamino acids (light and dark green), or (**C**) *S. enterica* cell cultures at different sampling points over time along rich (purple) and poor (red) nutrient growth curves. We found that the noise in cell length was larger than the noise in cell width.

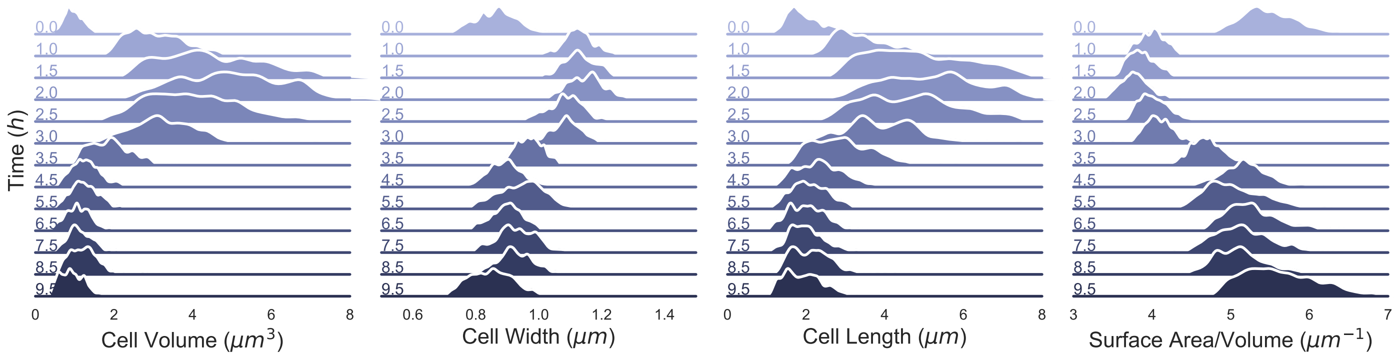


Figure S3: **Distributions of cell volume, width, length and surface-to-volume ratio for *E. coli* in nutrient-limited LB.** Full distributions of cell volume, width, length and surface-to-volume ratio estimated from segmented cells in microscopy images are shown along a growth curve in nutrient-limited LB. $t$ = 0 at the top indicates the overnight sample and subsequent samples are shown on the vertical axis. We found a broadening of the cell volume and length distributions around the time of the peak (1.5-3h). Cell width distributions did not become broader but the average cell width shifted earlier than that of the cell length.

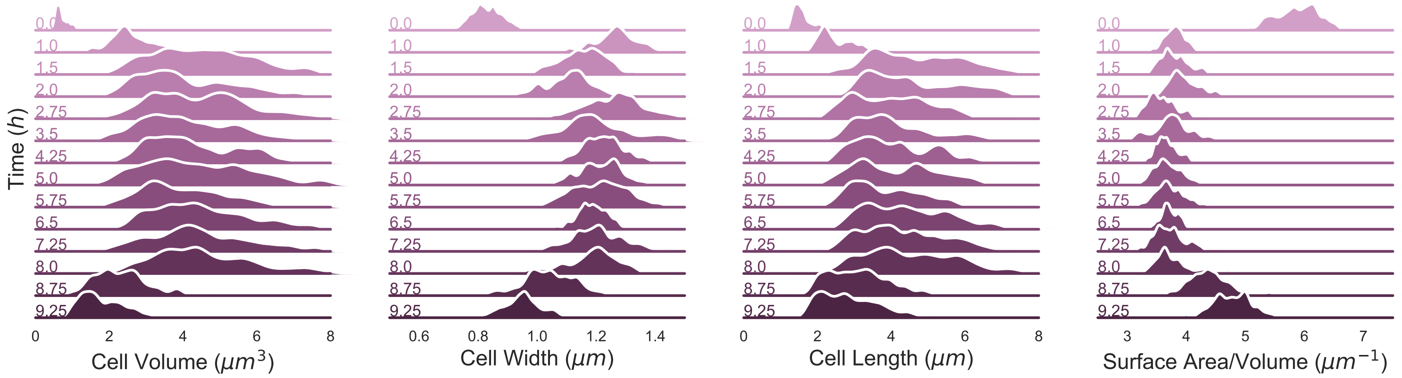


Figure S4: **Distributions of cell volume, width, length and surface-to-volume ratio for *E. coli* in sustained exponential growth in LB.** Histograms of cell volume, width, length and surface-to-volume ratio estimated from segmented cells in microscopy images are shown for sustained exponential growth in LB. $t = 0$ at the top indicates the overnight sample and subsequent samples are shown on the vertical axis. Sustaining exponential growth in LB media preserved the cell volume, length and width distributions. Surprisingly, the broadness of the cell volume and length distributions was also preserved, indicating that the noise in cell size was not caused by a transition from poor to rich medium.

### S1 Stochastic model for division control

In the main article, we used the mean-field approach to understand how mean cell size dynamics arise from the synchronization of the two processes, cell growth and division. However, our experiments reveal significant variation in cell volume, with the extent of this variation changing along the growth curve. This heterogeneity in cell volume plays a critical role in bacterial adaptability and survival, potentially enhancing stress tolerance and contributing to antimicrobial resistance [17, 41]. Understanding the sources of noise in cell volume is therefore key to deciphering the mechanisms of cell volume regulation [44] and for predicting bacterial population behaviour.

Here, we extend our mean-field approach to study noise in cell size by using an agent-based algorithm, where each cell has individual attributes and, therefore, individual cell size dynamics [10]. During cell growth, cells elongate exponentially at a *fractional* rate $\mu(t)$, therefore following the continuous differential equation:

$$\frac{ds}{dt} = \mu(t)s. \tag{S1}$$

In the main text, we describe that the rate of cell division depends on the growth rate, the cell size, and the division coefficient, i.e., $\mu(t)k(t)s$. In the stochastic model, the division control will be represented by an internal variable $x$, which can be interpreted as the level of a cell cycle regulator and follows the stochastic differential equation:

$$dx = \underbrace{\mu(t)k(t)sdt}_{\text{Deterministic dynamics}} + \underbrace{\sigma(s)dW}_{\text{Noise}}, \tag{S2}$$

where the first part of the equation represents the deterministic accumulation of the cell cycle regulator and the second part describes stochastic fluctuations. $W(t)$, corresponds to the Wiener process, which satisfies $\langle W(t)\rangle = 0$, and $\langle W(t)W(t')\rangle = \min(t, t')$ and is often employed to describe the levels of molecules subjected to random perturbations, including chemical compounding, synthesis, and degradation. $\sigma(s)$ modulates the noise and follows the expression:

$$\sigma(s) = \sigma_0 \left(\sqrt{\mu(t)k(t)s}\right) s^{\alpha}, \tag{S3}$$

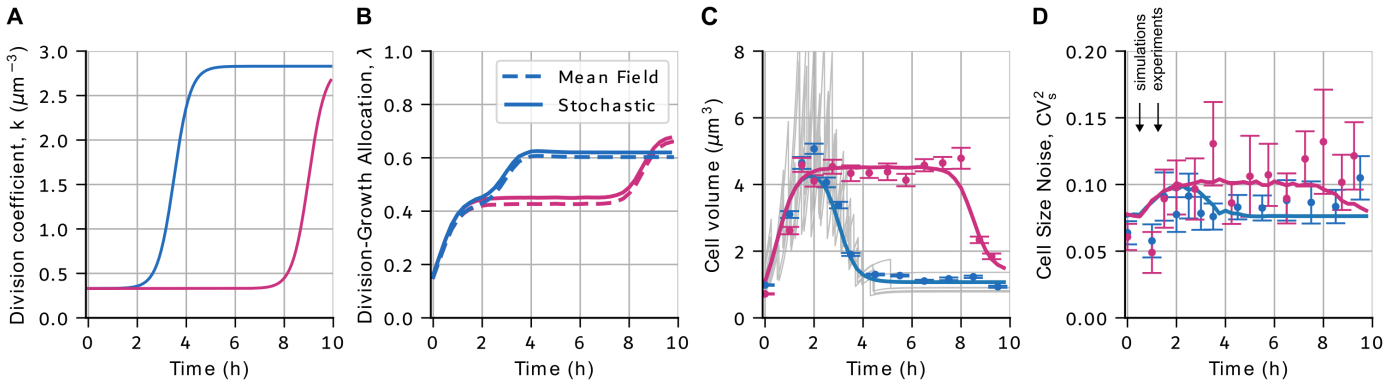


Figure S5: **A stochastic division model reproduces the noise in cell size dynamics from experiments with *E. coli* in LB. (A)** Dynamics of the division rate $k$ in LB along the growth curve (blue) and for sustained exponential growth (pink) using a stochastic cell division model. **(B)** Dynamics of the division-growth allocation parameter $\lambda$ over the growth curve from the mean-field approximation (dashed line; see also Fig. 4D) and stochastic cell size simulations (solid line). **(C)** Mean cell volume predictions from stochastic simulations (solid line) compared to experimental data (dots with error bars). Examples of stochastic cell size trajectories are shown in grey. **(D)** Noise ($CV_s^2$) of cell size over the growth curve or for sustained exponential growth from stochastic simulations (solid line) and experimental data (dots with error bars). Arrows indicate the increase in noise after resuspension ($t = 0$). Simulation dynamics were estimated over five thousand different cells for each time point tracking only one cell after division. The 95% confidence interval of the simulations is similar to the line width. Cell size noise was calculated by simulating the stochastic differential equation (S2) and fluctuations given by (S3) with $\sigma_0 = 0.23$ and $\alpha = 0.3$. At cell division, cell size is multiplied by a random variable $\beta$, which is beta-distributed so that $\langle\beta\rangle = 0.5$, and $CV_\beta^2 = 0.016$.

where $\sigma_0$ is a fitting constant, the term inside the parentheses is, except for the $dt$, exactly the term for the deterministic dynamics in the left of (S2) and the exponent $\alpha > 0$ defines a power law function of cell size. Depending on $\alpha$, cell size noise will be more or less tied to the mean cell size dynamics.

Cell division is triggered when the cell cycle regulator reaches a critical level $x = 1$ (for our units). At division, the cell size $s$ and the regulator $x$ undergo the following resets:

$$\text{if } x \geq 1 : \begin{cases} s \rightarrow s\beta;\, \beta \sim Beta(a, b) \\ x \rightarrow 0, \end{cases} \tag{S4}$$

where $\beta$ is a random variable that follows a beta distribution with parameters $a$ and $b$ such that the distribution has moments $\langle\beta\rangle = 0.5$ and noise $CV_\beta^2 = \frac{1}{2a+1}$, with $a = b$ being the shape parameters. We use a beta distribution since its support set is $(0, 1)$ and can be fully characterized using the first two moments (the mean and the noise).

Using the stochastic model to simulate the division, we fit the coefficient $k$ and, therefore, the division growth allocation $\lambda$ fitting the dependence of the noise modulator $\sigma$ with the mean cell size. However, the results on $\langle s\rangle$ are independent of $\sigma$ since this only affects the noise dynamics. The results of the mean cell size dynamics for LB yield similar results to the mean-field approach, which assumes that $\langle s^2\rangle \approx \langle s\rangle^2$ (Fig. 4D,E; S5A,B). The stochastic model predicts a consistently higher $\lambda$, meaning that the mean-field approach slightly underestimates the resources invested in division.

Similarly, the simulated mean cell size resembles the mean-field approximation very closely (Fig. 4C, S5C), but the stochastic model allows us now to also estimate the noise in cell size dynamics (Fig. S5D). To fit the cell size noise to our experimental observations, we made the noise modulation term in Equation (S2) to a function of $s$. In the simplest case, where the extrinsic noise is constant $\sigma(s) = \sigma$, the cell size noise $CV_s^2$ is also constant along the growth curve. However, our experiments suggest that cell size noise is higher for larger cell sizes and remains higher for sustained exponential growth (Fig. S3, S4). Therefore, we consider a weak dependence of $\sigma$ on cell size using $\sigma \sim s^{0.3}$. This weak dependence on cell size predicts the higher sustained noise during sustained exponential growth (Fig. S5D, pink) as well as the peak in noise along the growth curve (Fig. S5D, blue). However, the stochastic simulations predict an increase in the cell size noise slightly earlier than it occurs in the experiments (1h in the simulations versus 1.5-2h in the experiments). This difference could suggest a delay in the noise response or additional sources of noise that we are not accounting for. Noise might, for example, differ along the growth curve due to the expression of different stress factors, which could influence noise. Investigating factors that influence cell size noise could provide interesting insight into the effect of gene expression profiles along the growth curve on cell size control.
